# Overcoming preservation challenges to enable single-cell proteomics of fixed cell and tissue samples with retained proteome integrity

**DOI:** 10.1101/2025.03.10.642380

**Authors:** Agata N. Makar, Jocelyn Holkham, Sergio Lilla, Simon Wilkinson, Alex von Kriegsheim

## Abstract

The ability to assay the molecular composition of biological systems with single-cell resolution has revolutionised our understanding of tissue heterogeneity and function. Recent advances in single-cell proteomics (SCP) now enable the unbiased quantification of the proteome to a depth of several thousand proteins across hundreds of cells. Yet the broader adoption beyond specialised groups remains limited due to the need for specific equipment and expertise. A major challenge in making these analyses more broadly available is sample preservation for transporting biological material to SCP-capable facilities. To address this issue and provide practical solutions; we first evaluated various cell preservation methods from monolayer culture samples, then tested our optimised methodology on both cultured cells and, for the first time, preserved animal tissue from an *in vivo* mouse model. Our findings highlight the feasibility of SCP analyses in preserved tissues, significantly expanding its current applicability. By optimising upstream processing, our approach enables robust single-cell proteome analysis of both cells and tissues, making SCP more accessible to the wider scientific community. Ultimately, this advancement expands the potential applications of SCP, particularly in disciplines where analysing rare or heterogeneous populations is beneficial.

## Introduction

Single-cell proteomics (SCP) has emerged as the most comprehensive method to study cellular heterogeneity. It enables to distinguish and investigate diverse cell types and states, as well as their responses to genetic modifications, drug treatments, and other stimuli at the individual cell level^1,2^. Unlike bulk proteomics, which averages signals across populations of cells, SCP preserves single-cell resolution, revealing intricate biological details that are often obscured in the traditional bulk analyses^3,4^. This level of detail is critical for understanding diverse cellular dynamics, particularly in heterogeneous tissues, where distinct cell populations can drive disease progression or treatment resistance^5–8^.

Until the recent advancements in proteomics, transcriptomics was the primary method for analysing cells at single-cell resolution^9–11^. However, transcriptomic analyses are not able to provide information on post-translational regulation and modifications, which play essential roles in protein homeostasis and cellular function^12,13^. Consequently, transcript levels often show a poor correlation with protein abundance, partially due to dynamic regulation processes such as protein degradation and secretion^14,15^. Utilising SCP allows us to address these limitations by providing a direct readout of protein levels and enabling a more true representation of cellular states and functional changes.

Despite its transformative potential, SCP remains a technically complex and evolving field. Advances in the field have been driven by continuous optimisation of multiple steps such as cell isolation, processing, data acquisition methods, and computational analysis approaches^16–18^. With the development of highly sensitive mass spectrometers capable of detecting low-abundance protein inputs, one of the major challenges now lies in upstream sample processing. If not optimised, processing will lead to sample losses and reduced data depth resulting in significant changes in the proteome analysis depth and skew data interpretation by introducing artefacts that do not reflect the true physiological state of the cell. Sample preservation methods, buffer compositions, and storage conditions can impact cell integrity, proteome composition and peptide recovery, affecting data quality and reproducibility^19–21^. Cell and tissue samples often need to be transported between research laboratories and core facilities for processing and analyses, variations in preservation and handling can introduce inconsistencies. While various research groups have worked to refine workflows^22–24^, the challenge remains to identify a processing method that minimises proteome perturbations and preserves the biological state of cell and tissue samples as accurately as possible.

In this study, we comprehensively compared sample preservation methods for SCP. We examined monolayer cell samples in SCP format as well as extended this to challenging pancreatic tissue samples, to assess the impact of upstream processing. By systematically simulating shipping conditions through a 24-hour incubation at low temperatures, we identify the most effective preservation strategies for maintaining proteomic integrity while highlighting potential sources of data loss. Furthermore, we characterised specific classes of proteins that are particularly susceptible to loss during the aforementioned sample processing, which should be taken under consideration for specific research objectives.

Our findings establish a framework for best practices in SCP sample handling, aiming to improve the reliability of SCP analyses for the scientific community and making high-quality analyses available to laboratories that may not have direct, local access to the required instrumentation and expertise.

## Results

### Optimisation of sample preparation and data acquisition in single-cell proteomics

To ensure that our sample preparation workflow, including buffer composition, lysis conditions, digestion parameters, and data acquisition strategies, provided reliable and reproducible data, we first evaluated these parameters using a human colorectal carcinoma (RKO) cell line, comparing control (sgControl) and GSTP1 knockout (sgGSTP1) conditions. The knockout was introduced via CRISPR/Cas9 and not subcloned, therefore, we expected heterogeneity in GSTP1 expression levels stemming from a bi-allelic knockout to wild-type alleles.

Single-cell isolation and subsequent proteomic processing were performed following a standardised workflow (Fig. 1a). Briefly, cell suspensions were prepared in tissue culture, and individual cells were isolated using the CellenONE platform where lysis and digestion were also performed. Samples were subsequently loaded onto an Evosep One liquid chromatography (LC) system coupled with a timsTOF SCP mass spectrometer operating in parallel accumulation-serial fragmentation data-independent acquisition (diaPASEF) mode^25^.

**Figure 1.**
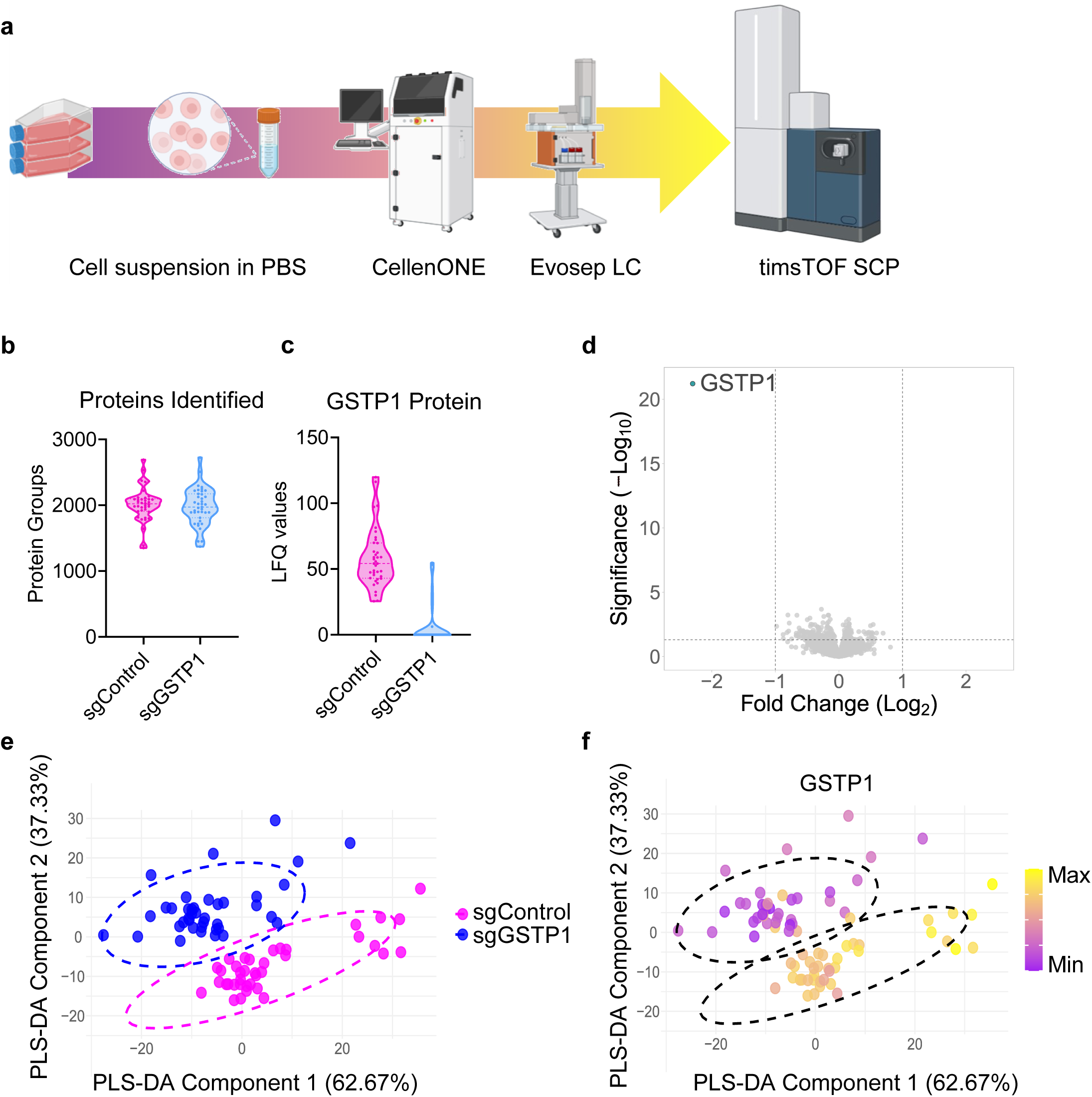
Optimisation of sample preparation and data acquisition in single-cell proteomics. (a) Schematic overview of the single-cell workflow. Created with Biorender. (b) Violin plot showing protein group identifications from single-cells in sgControl (magenta) and sgGSTP1 (blue). (c) MS analyses of GSTP1 levels in single-cell population. GSTP1 intensity is shown as label-free quantitation (LFQ) values. Each dot represents the readout from a single cell. (d) Volcano plot of two-sample *t-*test results (sgGSTP1 over sgControl). Log_2_ fold change and −log_10_ p-value are shown, significantly regulated proteins are shown in cyan. (e) PLS-DA plot of sample groups sgControl (blue) and sgGSTP1 (magenta). (f) PLS-DA plot from (e) overlaid with absolute values of GSTP1. The lowest GSTP1 value in the dataset is represented in purple (Min), the highest GSTP1 value in the dataset is represented in yellow (Max). sgControl n=38, sgGSTP1 n=39.

Cell lysis was performed using a buffer containing N-dodecyl β-D-maltoside (DDM), a mild detergent previously shown to be suitable for single-cell proteomics ^26^. The acquired proteomic data demonstrated a high degree of reproducibility, with a consistent number of proteins detected across single cells (Fig. 1b). Notably, we observed slight variations in GSTP1 expression levels among individual cells of the same genotype, underscoring the utility of single-cell analysis in capturing cellular heterogeneity (Fig. 1c). However, a distinct difference in GSTP1 levels between sgControl and sgGSTP1 samples was evident (Fig. 1d). Partial least squares discriminant analysis (PLS-DA) revealed clear separation between the two groups (Fig. 1e), further supporting the fidelity of our workflow. The absolute GSTP1 expression values corroborated this clustering pattern (Fig. 1f). We were able to quantify GSTP1 in some cells derived from the knockout cohort. The expression levels were lower than in the control cohort, suggesting that these cells were heterozygote for the wild-type allele. To further assess the quality of the data analyses, we utilised an entrapment sequence method using an *E.coli* library^27^. To achieve this, we extended the target human database to include the entrapment *E.coli* database, which contained protein sequences known to be false for our dataset. The percentage of shared proteins remained respectively low for all three FDR settings (Supplementary Fig. 1), giving us confidence in the search engines and the analysis pipeline.

Overall, these findings confirm that our lysis, digestion, data acquisition and analysis protocols are robust and reliable, supporting their application in further studies aimed at optimising upstream sample preservation strategies.

### The impact of freezing on single-cell proteomic profiles

Preservation strategies are crucial for maintaining sample integrity when immediate processing is not feasible and are essential to broaden the SCP user base. Given the high level of technical expertise and equipment needed to implement a sensitive SCP pipeline, the transport of preserved cells is a crucial element in this democratisation. To evaluate the effects of transport and cryopreservation on the single-cell proteome, we froze cells in either 10% or 20% dimethyl sulfoxide (DMSO) in foetal bovine serum (FBS) (Fig. 2a). Following 24 hours of storage at −80°C, cells were washed to remove cryoprotectant and processed according to the aforementioned workflow, using fresh samples as a benchmark.

**Figure 2.**
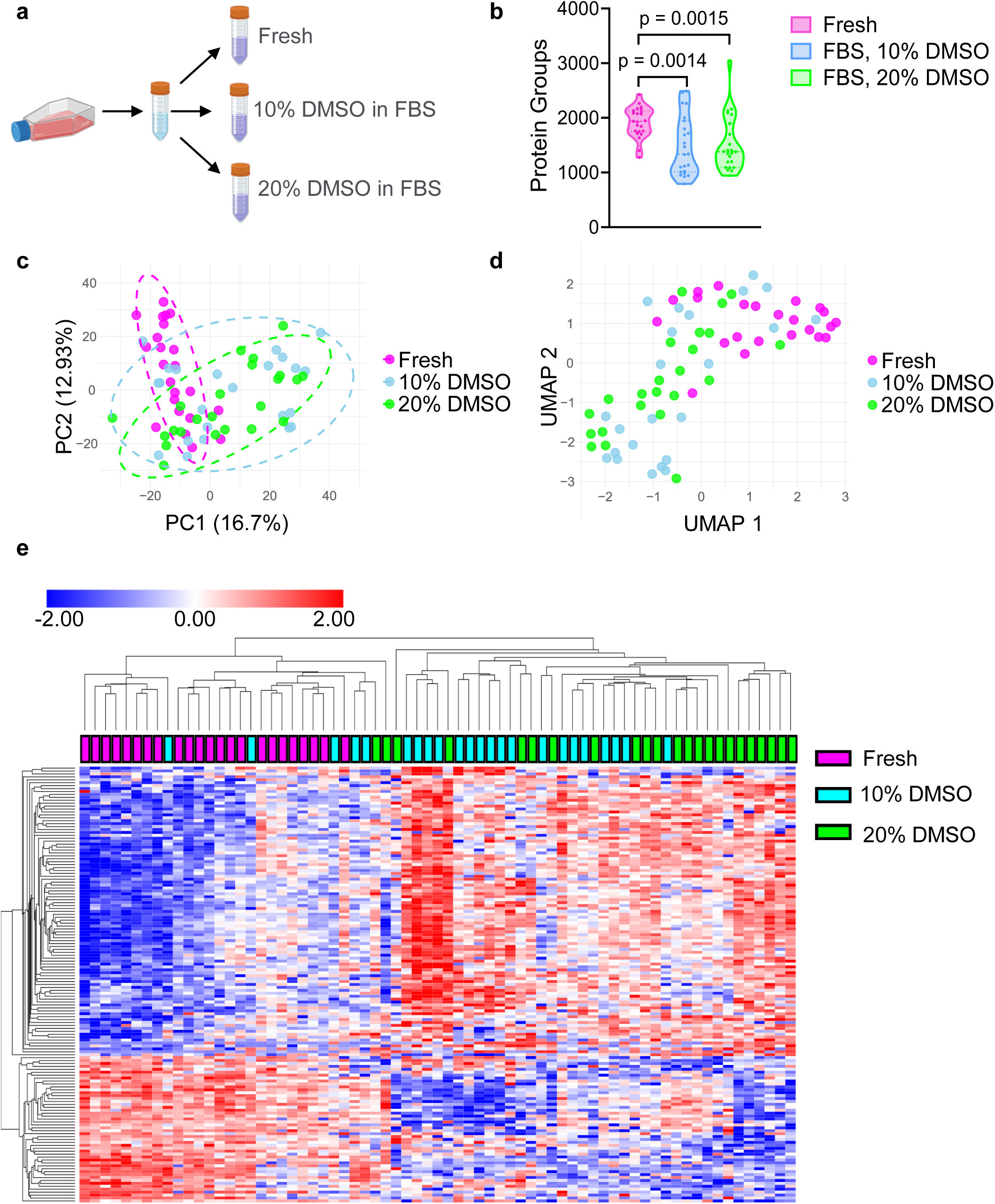
The impact of freezing on single-cell proteomic profiles. (a) Schematic overview of the single-cell cryopreservation workflow. Created with Biorender. (b) Violin plot showing protein group identifications from single-cells in tested conditions. Fresh (magenta), 10% DMSO in FBS (blue), 20% DMSO in FBS (green). Each dot represents the readout from a single-cell. P-values were calculated using an unpaired, two-tailed t-test. (c) PCA plot of sample groups as seen in (b) (d) UMAP of sample groups as seen in (b) (e) Heatmap of top 150 variable proteins in the tested dataset. P-values were calculated using one-way ANOVA. Hierarchal clustering of columns (samples) and rows (proteins detected) was performed using Euclidean distance. Fresh n=23, 10% DMSO n=23, 20% DMSO n=23.

Freezing and subsequent thawing led to a significant reduction in the number of proteins detected by mass spectrometry compared to fresh samples (Fig. 2b). Principal component analysis (PCA) revealed a clear distinction between frozen and fresh samples, with greater overlap observed among frozen conditions (Fig. 2c). Given the superior performance of uniform manifold approximation and projection (UMAP) over PCA for analysing large single-cell datasets^28^, we employed UMAP to visualise sample clustering. The results again demonstrated clear separation between fresh and frozen samples, with greater divergence observed in samples preserved with 20% DMSO (Fig. 2d). These findings were further corroborated by hierarchical clustering of the most variable proteins, where fresh samples formed a distinct cluster separate from frozen samples, with the greatest disparity observed between fresh and 20% DMSO conditions (Fig. 2e).

To pinpoint specific proteomic alterations induced by cryopreservation, we conducted differential expression analysis using volcano plots, revealing significant changes in protein abundance between fresh and frozen samples (Supplementary Fig. 2a, b). These observations align with previous reports demonstrating extensive proteomic alterations following cryopreservation^22,23^. Moreover, the requirement for ultra-low temperature storage and transport of frozen samples imposes additional logistical and financial constraints on experimental workflows.

Thus, our results indicate that cryopreservation is not a recommended method for preserving single cells for proteomic analysis due to significant alterations and the logistical and financial challenges associated with ultra-low temperature storage and transport.

### Minimal paraformaldehyde fixation for single-cell proteomics preserves proteome integrity

To improve sample preservation in single-cell proteomics, we evaluated paraformaldehyde (PFA), a polymeric form of formaldehyde widely used for cell and tissue fixation^29^. A recent study demonstrated the feasibility of formaldehyde fixation for single-cell proteomics^24^. However, we sought to minimise fixative concentration while maintaining proteomic integrity, therefore, we tested 0.1%, 0.3%, and 1% PFA in PBS, with fresh cells serving as a benchmark.

Given the low PFA concentrations used, we extended the fixation time to 24 hours before single-cell sorting and processing. Quantification of protein identifications across conditions showed no decrease in detected proteins compared to fresh cells, in contrast to cryopreserved samples (Fig. 3a).

**Figure 3.**
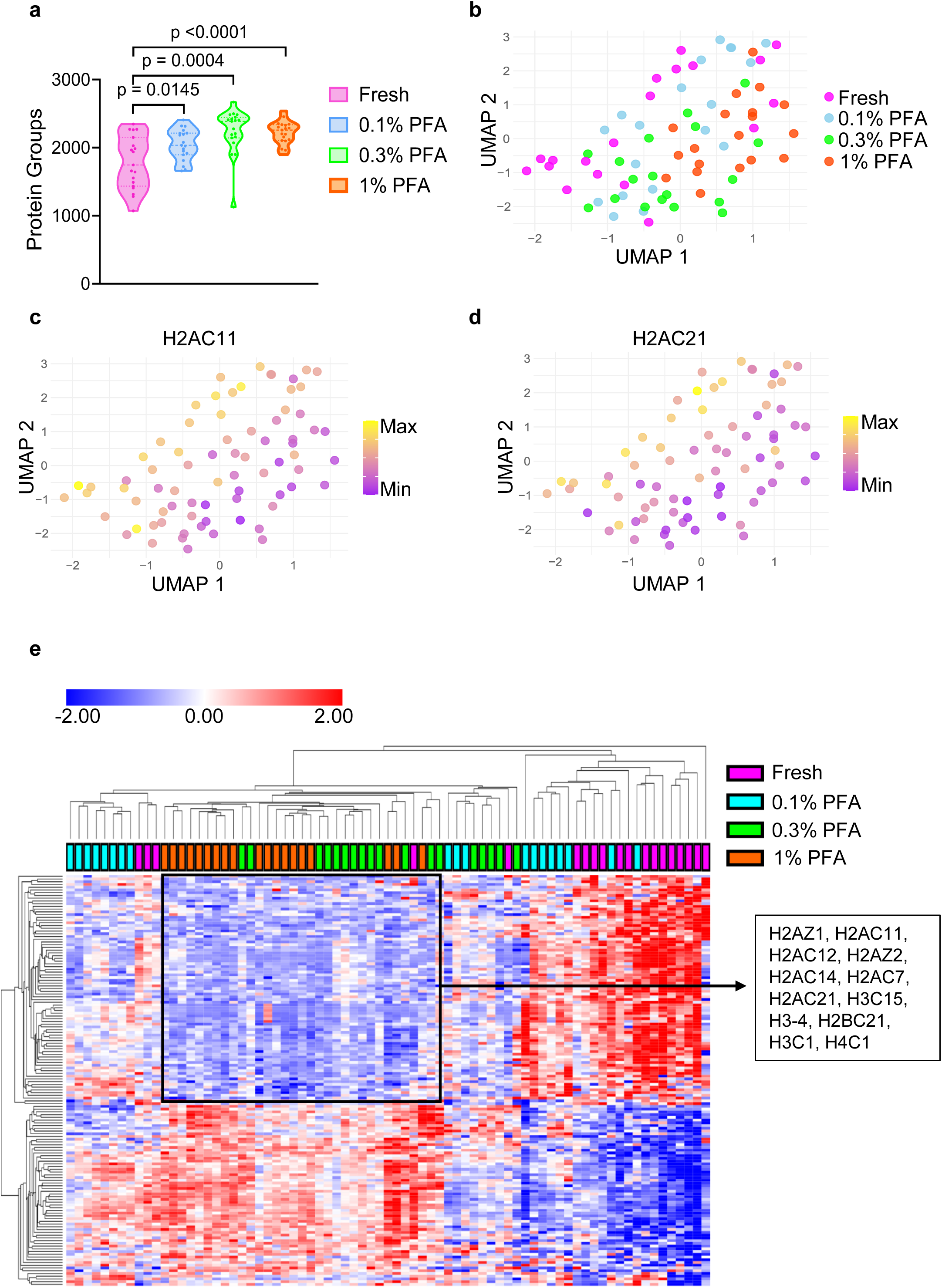
Minimising paraformaldehyde fixation for single-cell proteomics preserves proteome integrity. (a) Violin plot showing protein group identifications from single cells in tested conditions. Fresh (magenta), 0.1% PFA (blue), 0.3% PFA (green), 1% PFA (orange). Each dot represents the readout from a single-cell. P-values were calculated using an unpaired, two-tailed t-test. (b) UMAP of sample groups as seen in (a) (c) UMAP plot from (b) overlaid with absolute values of H2AC11. The lowest H2AC11 value in the dataset is represented in purple (Min), the highest H2AC11 value in the dataset is represented in yellow (Max). (d) UMAP plot from (b) overlaid with absolute values of H2AC21. The lowest H2AC21 value in the dataset is represented in purple (Min), the highest H2AC21 value in the dataset is represented in yellow (Max). (e) Heatmap of top 150 variable proteins in the tested dataset. P-values were calculated using one-way ANOVA. Hierarchal clustering of columns (samples) and rows (proteins detected) was performed using Euclidean distance. Selected histone proteins from the highlighted cluster are listed on the right. Fresh n=19, 0.1% PFA n=19, 0.3% PFA n=19, 1% PFA n=19.

UMAP analysis revealed distinct clustering patterns, with fresh and 0.1% PFA-fixed cells grouping in the top left quadrant and 0.3% and 1% PFA-fixed cells clustering in the opposite direction (Fig. 3b). This suggests that 0.1% PFA fixation best preserves the native proteome. Comparison of each fixation condition to fresh cells using volcano plots indicated a progressive increase in significantly reduced (i.e., lost) proteins with increasing PFA concentration (Supplementary Fig. 3a–c).

Further analysis of proteins significantly depleted in the 1% PFA condition revealed a distinct cluster of chromatin-associated proteins (Supplementary Fig. 3d). Examination of absolute histone protein levels in UMAP space demonstrated a clear downward trend in abundance with increasing PFA concentration (Fig. 3c, d). Heatmap analysis of the most variable proteins further supported these alterations, with fresh and 0.1% PFA-fixed cells clustering together and an enrichment of histone proteins in these groups compared to 0.3% and 1% PFA (Fig. 3e). The depletion of histone proteins at higher PFA concentrations is likely due to extensive cross-linking of lysine residues, reducing peptide digestion efficiency^30^.

Interestingly, the same heatmap cluster also contained multiple exosome-associated proteins, suggesting increased secretion in cells fixed with 0.3% and 1% PFA (Supplementary Fig. 3d). Indicating that higher PFA concentrations may introduce artefacts, making them unsuitable for certain experiments.

Overall, our results demonstrate that 24-hour fixation with low PFA concentrations preserves proteomic integrity. Fixation with 0.1% PFA provides results comparable to fresh cells, whereas higher PFA concentrations induce significant proteomic changes and should be avoided.

### Short-term fixation is sufficient to preserve proteome integrity

To further minimise the impact of sample preparation on single-cell proteomics data, we evaluated the effects of a shorter fixation duration. Cells were fixed for 20 minutes with PFA, followed by quenching with tris-buffered saline (TBS). Post-fixation, cells were washed to remove residual fixative, incubated at 4°C for 24 hours, and subsequently processed for analysis. Proteomic profiles from these fixed cells were compared to those of freshly processed cells. The number of proteins detected per cell was comparable across all fixation conditions (Fig. 4a). UMAP analysis of protein groups revealed no distinct separation between the conditions, although minor subclusters of fresh and 1% PFA-fixed cells were observed at opposite ends of the UMAP space (Fig. 4b).

**Figure 4.**
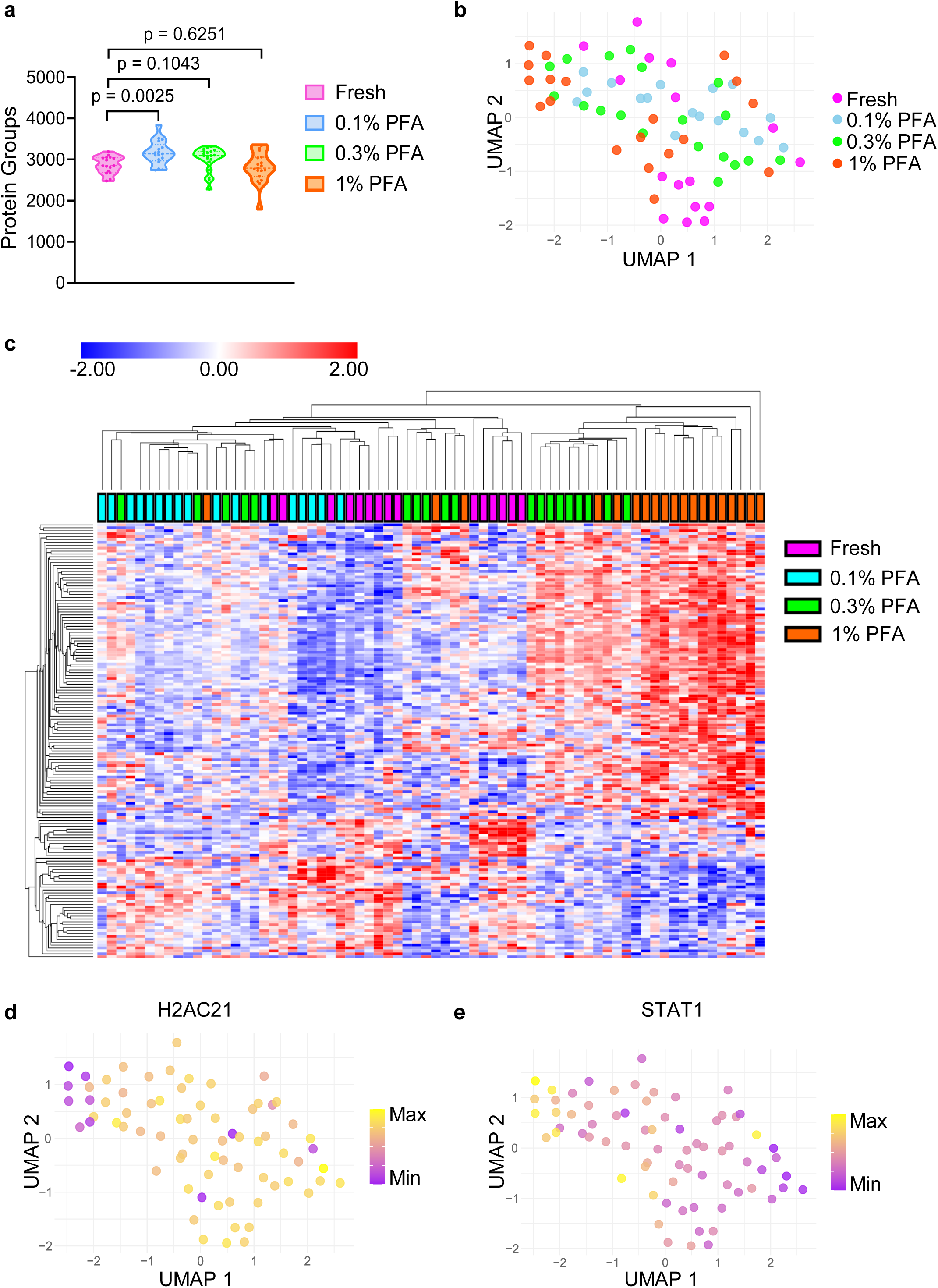
Short-term fixation is sufficient to preserve proteome integrity. (a) Violin plot showing protein group identifications from single cells in tested conditions. Fresh (magenta), 0.1% PFA (blue), 0.3% PFA (green), 1% PFA (orange). Each dot represents the readout from a single cell. P-values were calculated using an unpaired, two-tailed t-test. (b) UMAP of sample groups as seen in (a) (c) Heatmap of top 200 variable proteins in the tested dataset. P-values were calculated using one-way ANOVA. Hierarchal clustering of columns (samples) and rows (proteins detected) was performed using Euclidean distance. (d) UMAP plot from (b) overlaid with absolute values of H2AC21. The lowest H2AC21 value in the dataset is represented in purple (Min), the highest H2AC21 value in the dataset is represented in yellow (Max). (e) UMAP plot from (b) overlaid with absolute values of STAT1. The lowest STAT1 value in the dataset is represented in purple (Min), the highest STAT1 value in the dataset is represented in yellow (Max). Fresh n=16, 0.1% PFA n=17, 0.3% PFA n=19, 1% PFA n=19.

Volcano plot comparisons between each fixation condition and fresh cells demonstrated a progressive increase in significantly downregulated proteins with increasing PFA concentration, consistent with trends observed in 24-hour fixation (Supplementary Fig. 4a–c). Hierarchical clustering of the top variable proteins across conditions showed a distinct segregation of the 1% PFA and 0.3% PFA groups from the remaining samples (Fig. 4c). When examining histone proteins, which exhibited significant reductions at higher PFA concentrations in long-term fixation, no clear trend was observed in absolute values under short-term fixation. However, a cluster of 1% PFA-fixed cells still displayed a reduction in histone protein abundance (Fig. 4d). Interestingly, a subset of proteins, including STAT1, exhibited increased abundance at higher PFA concentrations, demonstrating an inverse correlation with histone protein levels (Fig. 4e).

Overall, these findings indicate that a 20-minute fixation duration is sufficient to preserve the proteome. However, 1% PFA should be avoided due to substantial proteomic alterations, whereas lower concentrations such as 0.1% PFA introduce minimal perturbations relative to fresh cells.

### Proteomic profiling of a polyclonal PDAC cell line

To extend our findings, we analysed a heterogenous, murine pancreatic ductal adenocarcinoma (PDAC) KPC (BL6J) cell line that was established by pooling cells from tumours formed within the KPC genetically-engineered mouse model of pancreatic cancer^31^, using 0.1% PFA for 20-minute fixation. This approach returned a consistent number of identified proteins across the cell population. UMAP analysis revealed a clear separation of the population into two distinct cell clusters (Fig. 5a), a pattern similarly observed in the PCA plot (Supplementary Fig. 5a). When comparing the number of proteins identified in these clusters, we noted a significant difference, with a smaller cluster (Cluster 2, shown in blue) exhibiting an increase in detected protein (Fig. 5b). Volcano plot comparisons of the two subpopulations, alongside heatmap analysis with hierarchical clustering of the top variable proteins, demonstrated distinct clustering with multiple proteins significantly enriched in Cluster 2 (Fig. 5c, Supplementary Fig. 5b). Gene ontology analyses (GO) of biological processes associated with these proteins revealed an enrichment of proteins involved in translation and biosynthetic processes (Fig. 5c, Supplementary Fig. 5c).

**Figure 5.**
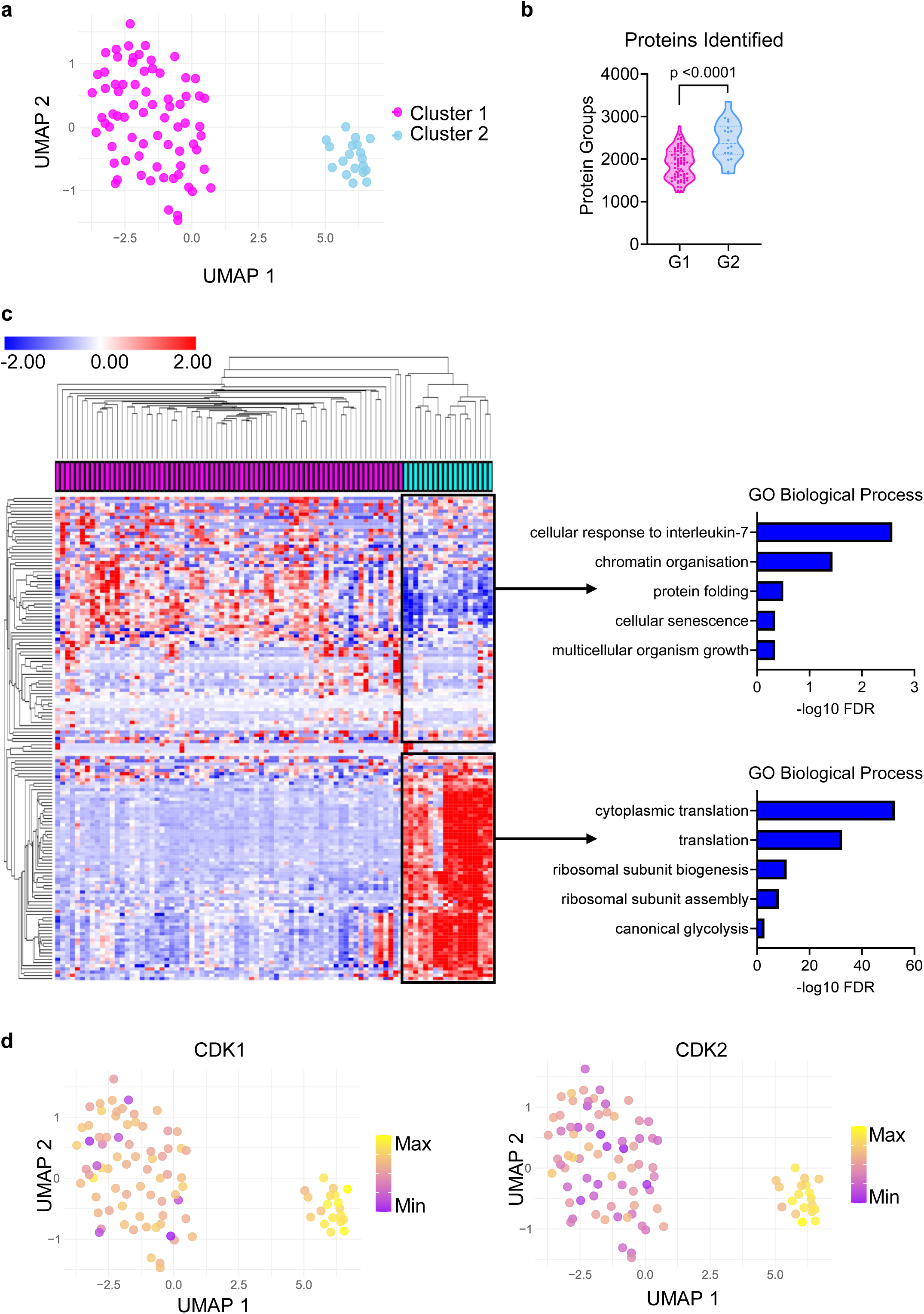
Proteomic profiling of a polyclonal PDAC cell line. (a) UMAP of sample groups Cluster 1 (magenta), Cluster 2 (blue). (b) Violin plot showing protein group identifications from single cells in two clusters. Cluster 1 (magenta), Cluster 2 (blue). Each dot represents the readout from a single cell. P-value was calculated using an unpaired, two-tailed t-test. (c) Heatmap of top 200 variable proteins in the tested dataset. Hierarchal clustering of columns (samples) and rows (proteins detected) was performed using Euclidean distance. Top 5 GO biological processes from main protein clusters are shown on the right. (d) UMAP plot from (a) overlaid with absolute values of CDK1 (left) and CDK2 (right). The lowest values in the dataset are represented in purple (Min), the highest values in the dataset are represented in yellow (Max). PDAC single-cell n=92.

Given this enrichment of translation-related proteins and the observed increase in protein detection within Cluster 2, we examined CDK1 and CDK2 levels. Both proteins play key roles in promoting cell cycle progression and their increased abundance suggests an active proliferative state^32,33^. Indeed, UMAP analysis of absolute CDK1 and CDK2 values confirmed their upregulation in Cluster 2 potentially explaining the observed phenotype (Fig. 5d).

Overall, these findings demonstrate that fixation for SCP is sufficiently robust to detect distinct cellular subpopulations.

### Proteomic alterations induced by cell preservation

A comprehensive comparison of various preservation methods revealed that while low concentrations of fixative (0.1% PFA) effectively preserve the cellular proteome and closely resemble the fresh cell state, none of the tested methods fully recapitulate the proteomic profile of fresh cells. To enable researchers across different disciplines to apply this methodology for cell population studies, we sought to identify protein groups sensitive to cryopreservation and fixation before single-cell sorting. This information is critical for experimental design and data interpretation.

To achieve this, we conducted GO analyses on biological processes and cellular components affected by upstream sample preparation across all tested cell cultures and conditions. As expected, GO analyses and clustering demonstrated distinct separation of cryopreserved samples, which exhibited significant enrichment in multiple biological processes and cellular components (Fig. 6a, b). These analyses also highlighted an enrichment of proteins involved in proteolysis regulation and metabolic processes across most conditions (Fig. 6a). Furthermore, a strong enrichment of secretory granule lumen proteins was observed, aligning with our findings of exosomal proteins loss during fixation (Fig. 3e). Among all tested conditions, 0.1% PFA fixation resulted in the lowest number of significantly reduced protein concentrations compared to fresh cells (data not shown). This trend was reflected in the lowest enrichment scores for the highlighted processes and cellular components in GO term heatmaps (Fig. 6a, b).

**Figure 6.**
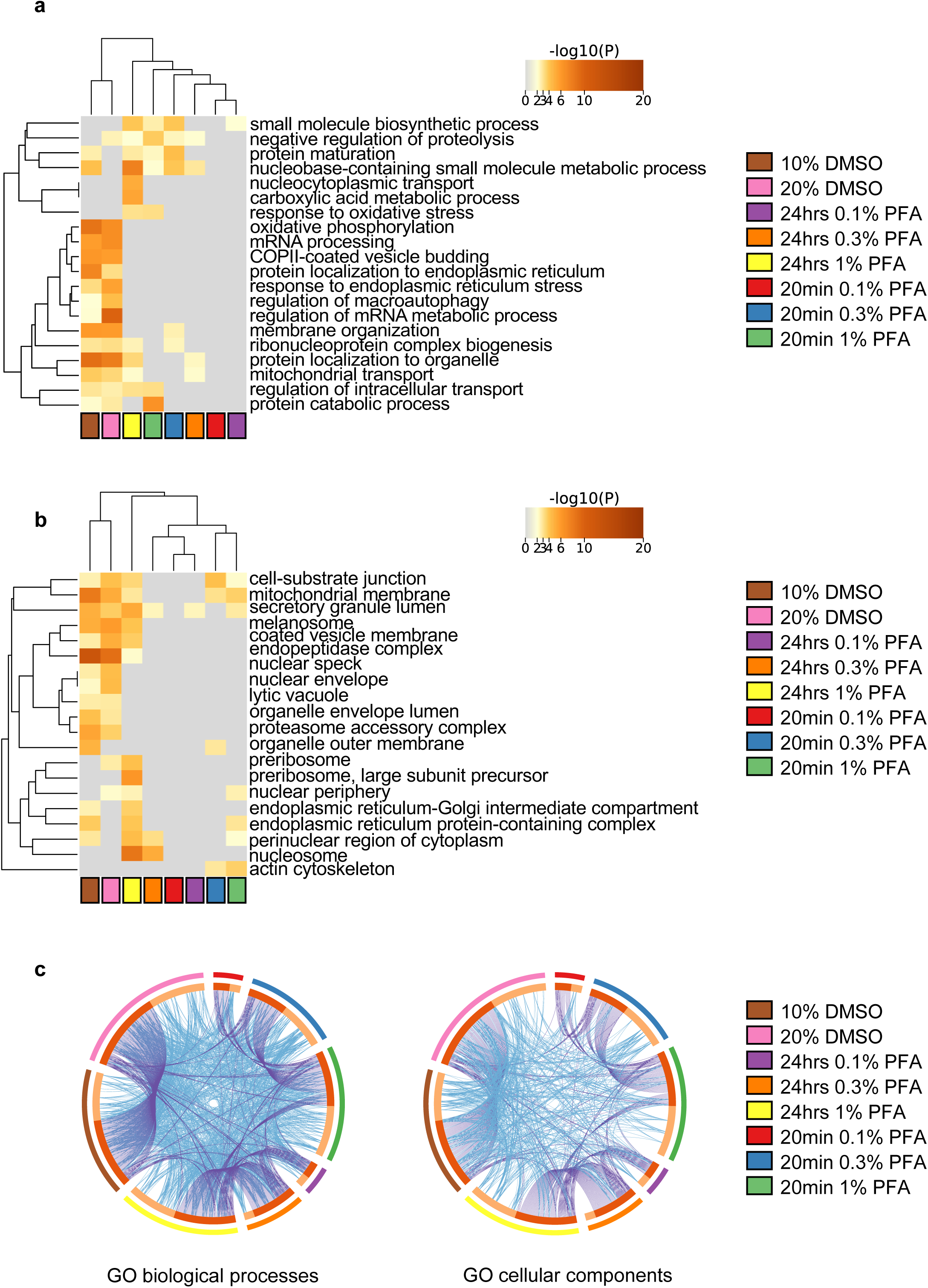
Proteomic alterations induced by cell preservation. (a) Functional clustering of the top 20 significantly enriched GO biological processes in the indicated datasets. The heatmap cells are coloured by their p-values with grey indicating a lack of enrichment for that term in the corresponding dataset. (b) Functional clustering of the top 20 significantly enriched GO cellular components in the indicated datasets, colour-coded as seen in (a). (c) Circos plot showing the proteome alterations induced by cell preservation shared between datasets. On the outside, each colour represents a different method tested and corresponds to the figure legend shown on the right. On the inside, the dark orange colour represents proteins that appear in multiple datasets and the light orange colour represents unique proteins specific to each tested condition. Purple lines represent proteins shared across the datasets. Blue lines represent proteins which fall into the same statistically significant GO term.

Circos plots^34^ depicting protein overlap across conditions (with overlapping proteins shown in purple) revealed intra-group similarities, such as overlap among different PFA concentrations following 24-hour fixation. However, only a limited number of proteins overlapped between different datasets (Fig. 6c). Overall, few proteins were commonly downregulated across conditions in terms of biological processes and cellular components. Notably, while the Circos plot demonstrated limited protein overlap, the associated biological processes and cellular components (represented in blue) exhibited a much higher degree of similarity between datasets (Fig. 6c). Collectively, these findings confirm that low PFA concentrations are a favourable approach for proteome preservation. Moreover, proteomic perturbations introduced by preservation methods primarily affect entire biological processes and cellular structures rather than individual proteins conserved across conditions.

### Single-cell proteomic profiling of fixed pancreatic tissue reveals distinct clusters

To evaluate the feasibility and effectiveness of low-level PFA fixation for preserving tissue samples for proteomic analyses, we focused on the mouse pancreas. This organ was chosen due to its defining feature of high protein production and secretion in acinar cells, which comprise the largest pancreatic cell population. Furthermore, the highly dynamic acinar cell population is often underrepresented in single-cell transcriptomic analyses^35,36^. It also presents a good technical test, given that acinar cells are an inherently autolytic compartment due to very high hydrolase content, these enzymes pose significant challenges for single-cell transcriptomic and proteomic analyses. We hypothesised that obtaining meaningful data from pancreatic acinar cells would demonstrate that our method is sufficiently robust to be implemented across even the most challenging mammalian tissues.

A pancreas from a wild-type BL6/129 mouse was dissociated into single cells (see *Materials and Methods*) and subjected to proteomic profiling. We adopted a 24-hour fixation period in 0.1% PFA, supplemented with protease inhibitors to mitigate autodigestion and preserve protein integrity^37,38^. Gating based on cell size allowed us to remove subpopulations of unwanted cells such as immune or endothelial cells allowing us to enrich the acinar cell population. Moreover, to ensure the downstream processing was restricted to intact cells we implemented DAPI staining step to exclude damaged, dead or dying cells where membrane integrity is compromised^39^. This optimised approach resulted in a consistent and reproducible number of proteins identified per cell across the dataset (Supplementary Fig. 6a).

Notably, amylase, which is a well-established marker of acinar cells, was consistently detected across all analysed cells, alongside several other key exocrine pancreatic enzymes (Fig. 7a). These findings are in agreement with data from the Human Transcriptome Cell Atlas, which similarly demonstrates the enrichment of digestive enzymes in the acinar cell population^40^. This enrichment of acinar cells seen in our proteomics and immunofluorescent (Supplementary Fig. 6b) analyses align with the known composition of pancreatic tissue, where acinar cells are the major cell type^41,42^.

**Figure 7.**
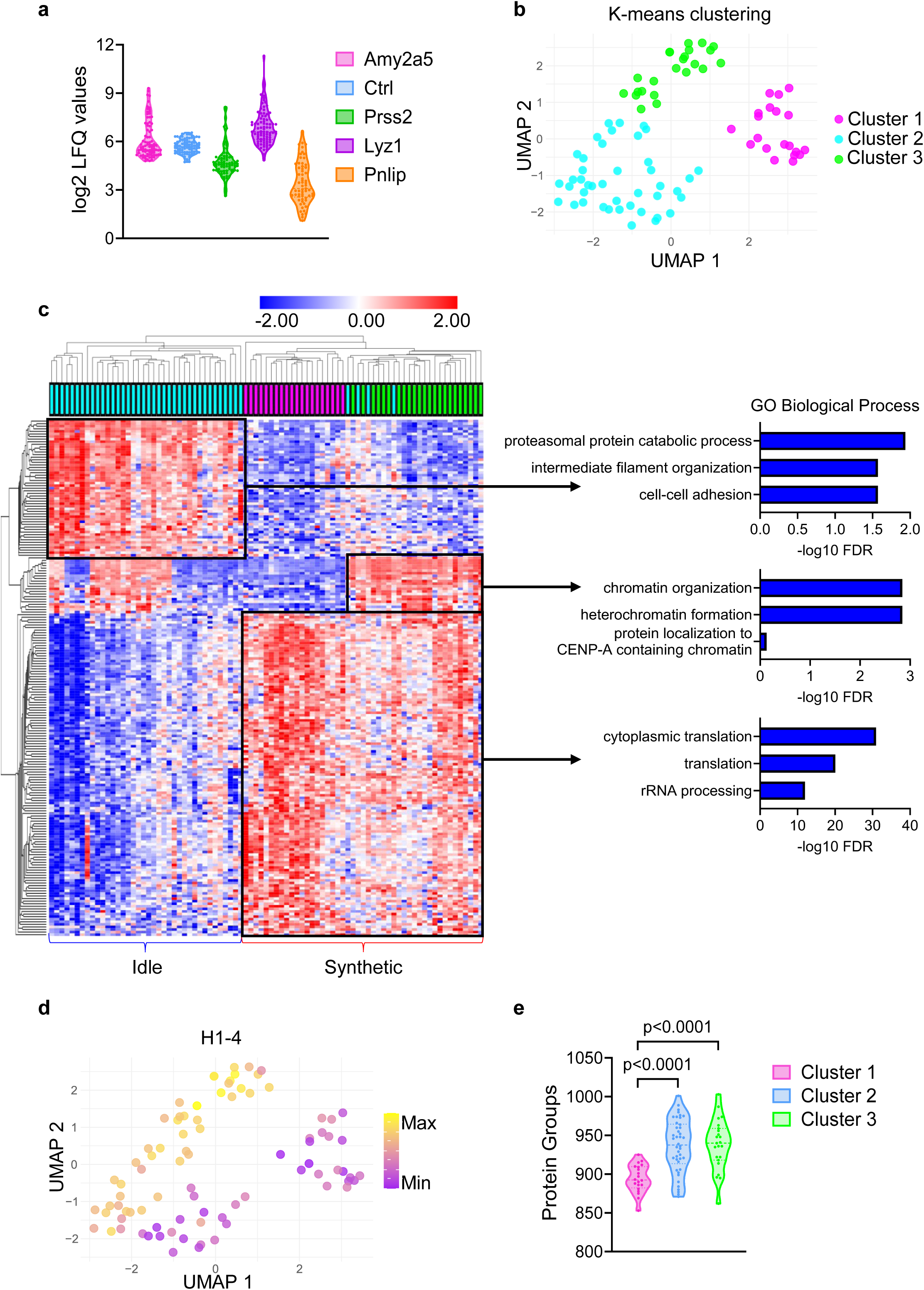
Single-cell proteomic profiling of fixed pancreatic tissue reveals distinct clusters. (a) Violin plot showing log2 LFQ values for selected pancreatic enzymes in single cells derived from pancreatic tissue sample. Each dot represents the readout from a single cell. (b) UMAP of sample groups separated based on k-means clustering (cluster number=3). Cluster 1 (magenta), Cluster 2 (blue), Cluster 3 (green). (c) Heatmap of top 200 variable proteins in the tested dataset based on p-values. P-values were calculated using one-way ANOVA. Hierarchal clustering of columns (samples) and rows (proteins detected) was performed using Euclidean distance. Top 3 GO biological processes from main protein clusters are shown on the right. (d) UMAP plot overlaid with absolute values of H1-4. The lowest H1-4 value in the dataset is represented in purple (Min), the highest H1-4 value in the dataset is represented in yellow (Max). (e) Violin plot showing protein group identifications from single-cells in three clusters. Cluster 1 (magenta), Cluster 2 (blue), Cluster 3 (green). Each dot represents readout from a single-cell. P-values were calculated using an unpaired, two-tailed t-test. Pancreas single-cell Cluster 1 n=20, Cluster 2 n=42, Cluster 3 n=23.

To further dissect the cellular heterogeneity within the acinar-enriched population, we employed k-means clustering on the single-cell proteomic profiles, stratifying the cells into three groups for further analyses (Fig. 7b). A heatmap of the most variable proteins among clusters further highlighted divergent expression patterns corresponding to the assigned groups (Fig. 7c). Comparative analysis of relative protein intensities across the clusters showed distinct expression patterns, underscoring the inherent heterogeneity within the acinar cell population (Supplementary Fig. 6c).

Analysis of the differential abundance of translational machinery across clusters revealed unique profiles of protein dynamics, enabling us to broadly categorise the cells into ‘synthetic’ (high translation) and ‘idle’ (low translation) acinar states (Fig. 7c). This observation is consistent with previous studies that have reported similar functional stratifications within the acinar cell population based on transcriptome data^43^. Importantly, the expression patterns of several pancreatic digestive enzymes varied between clusters and did not directly correlate with levels of translational machinery (Supplementary Fig. 6d). This suggests that although translation processes are upregulated at synthetic acinar state, in some subpopulations this does not directly correspond to protein levels within the cells, possibly due to post-translational events such as increased release of enzymes from the cells, which cannot be detected at the RNA level. It also highlights a key advantage of proteomic analyses over transcriptomics, especially in tissues with dynamic protein secretion, where protein-level information offers a more nuanced and accurate depiction of the cellular state.

A deeper investigation into the cell clusters revealed that cluster 1, which belongs to the ‘synthetic’ subpopulation, exhibited a significantly lower number of protein IDs compared to the other clusters. This reduction in protein IDs was correlated with decreased levels of histone proteins, such as H1-4 (Fig. 7d). Interestingly, visualisation of H1-4 levels using UMAP indicated that a subset of cells within cluster 2 (‘idle’ subpopulation) also displayed reduced abundance relative to the rest of the population. By further stratifying cluster 2 based on H1-4 levels (Supplementary Fig. 6e), we identified significant differences in protein IDs between the resulting subclusters (Supplementary Fig. 6f). These differences in histone protein abundance could indicate differences on genome level as tetraploid genomes are prevalent in mouse acinar cells^44^. Alternatively, these findings corroborate previous reports suggesting that histone turnover can serve as an indicator of proliferative cells and provide an additional parameter for distinguishing cellular subpopulations^45^.

Collectively, our results demonstrate that low-level PFA fixation is a robust and effective method for preserving proteomic integrity in pancreatic tissue, enabling the resolution of distinct cellular subpopulations. This approach provides critical insights into cell states and functional heterogeneity, particularly with regard to protein synthesis and secretion dynamics, which are often challenging to capture using transcriptomic techniques alone. These findings underscore the value of single-cell proteomics in revealing the complex biology of highly secretory tissues such as the pancreas.

## Discussion

The impact of sample preservation on single-cell proteomics workflows remains a critical yet underexplored factor influencing data quality and biological interpretation. Our study systematically assessed the effects of cryopreservation, PFA fixation, and fixation duration, while also mimicking shipping conditions. We observed distinct effects of these parameters on protein recovery, quantification, and overall proteomic profiles, revealing nuanced consequences of each method.

DMSO-based freezing is widely used as a preferred technique for cell preservation in single-cell RNA sequencing studies^46^. However, we did not observe a comparable advantage for SCP analyses. This finding aligns with previous reports demonstrating that even minimal DMSO exposure (e.g., 0.1%) induces significant alterations in the cellular proteome^47,48^. Moreover, cryopreservation can introduce biomechanical stress, further distorting the cellular proteome^22^. These findings raise important considerations about using DMSO-based cryopreservation for single-cell analyses, including RNA and protein studies. Given that cryopreservation can induce cellular stress and potential damage, it may influence the integrity and reliability of data obtained from these approaches, underscoring the need for further evaluation of its impact.

Our analysis demonstrated that low concentrations of PFA preserve the proteome in a state most reflective of native cellular conditions. However, careful optimisation is essential, as PFA fixation can introduce artefacts. We observed a significant reduction in histone levels with increasing PFA concentrations, likely due to the high lysine content of histones, which promotes excessive crosslinking and impairs trypsin digestion^49,50^. Incorporating chemical de-crosslinking steps prior to proteomic processing may improve protein recovery in highly cross-linked samples^51^, although such approaches could result in substantial data loss in single-cell settings due to low working volumes and input material. Alternatively, cellular stress responses, including enhanced vesicle secretion and histone release, may contribute to the observed depletion of histones at higher PFA concentrations as recently reported^52^. This interpretation is further supported by our data, which revealed a concomitant reduction of exosomal proteins with high-PFA conditions, suggesting a potential relationship between fixation and exosome-mediated histone secretion.

The intricate interplay between fixative choice and cellular proteome integrity must be considered, particularly in the study of highly dynamic and sensitive pathways, such as signalling cascades. An illustrative example from our dataset is the observed enrichment of STAT1 in samples fixed with higher PFA concentrations, in inverse correlation with histone protein levels. Given that STAT1 stability is regulated through ubiquitination at lysine residues^53^, it is plausible that fixation-induced crosslinking obstructs this regulatory mechanism, leading to an accumulation of STAT1. These findings underscore the necessity of accounting for fixation-induced alterations when designing SCP experiments and highlight the advantages of low-PFA fixation in mitigating such artefacts.

For the first time, we applied single-cell proteomics to fixed tissue samples, specifically pancreatic tissue, which presents unique challenges due to its high concentration of digestive enzymes. These enzymes contribute to RNA and protein degradation, complicating transcriptomics and proteomics analyses, respectively^54^. Additionally, the tissue’s secretory function, which leads to high secretion of newly expressed proteins, will amplify discrepancies between RNA and protein levels^55–57^. Our optimised preservation strategy enabled us to delineate distinct acinar cell subpopulations by minimising autodigestion and excluding background cells such as immune or necrotic populations. Interestingly, we identified a subset of cells exhibiting a proteomic signature indicative of high protein synthesis which did not directly correlate with pancreatic enzyme levels, histone protein abundance, or overall proteome content as measured by the number of detected protein IDs. These observations suggest the presence of previously unrecognised acinar subpopulations or subtle protein dynamic shifts that remain undetectable in transcriptomic studies. These findings have significant implications, not only for fundamental cell biology but also for future identification of cellular states susceptible to therapeutic interventions or markers indicative of pathological progression.

Overall, our study advances the field of single-cell proteomics by providing a comprehensive evaluation of cell and tissue preservation strategies, offering guidance for optimised SCP workflows that minimise alterations while preserving proteomic integrity.

## Materials and Methods

### Cell culture and treatment

RKO, MDA-MB-231 and PDAC cells were cultured in Dulbecco’s Modified Eagle’s Medium (DMEM) containing 4.5 g/L glucose, sodium bicarbonate, ʟ-glutamine and sodium pyruvate (Sigma-Aldrich, D6429) and supplemented with 10% (v/v) Foetal Bovine Serum (FBS) (Gibco, 16000044). All cells were maintained in a humidified chamber maintained at 37°C and 5% CO_2_.

### GSTP1 gene editing

crRNA-mediated knockout in RKO cells was performed by CRISPR/Cas9-mediated gene editing using Nucleofection. Predesigned Alt-R™ CRISPR-Cas9 guides RNA (IDT) targeting the following human sequences: Hs.Cas9.GSTP1.1.AA 5’GGGAAATAGACCACGGTGTA; Hs.Cas9.GSTP1.1.AB 5’ AATACCATCCTGCGTCACCT. Alt-R® CRISPR-Cas9 and a Negative Control crRNA #1 (IDT, 1072544) were used. To form Cas9-RNP complex, all crRNAs were annealed with Alt-R CRISPR-Cas9 tracrRNA (IDT, 1072533) at 95 °C before adding Cas9 (IDT, 1081058) at room temperature (RT). The Cas9-RNP complexes with Alt-R Cas9 Electroporation enhancer (IDT, 1075916) were transfected into cells using Amaxa SE Cell Line 4D-Nucleofector X Kit S (Lonza, V4XC-1032).

### Immunofluorescence staining

4 μm sections were cut from formalin-fixed, paraffin-embedded samples (FFPE) onto charged slides then placed in a 37°C oven overnight. Slides were de-waxed in xylene, then rehydrated with graded ethanol. Antigen retrieval was performed by immersing the slides in 10 mM sodium citrate, pH 6.0 and boiling them for 10 min using a microwave. Tissue was then permeabilised in 0.1% Triton X-100 in PBS for 10 min and blocked with protein block (5% goat serum, 0.3% Triton X-100 in PBS) for 30 min at RT. After three washes with 0.1% Triton-X-100 in PBS for 2 minutes, tissues were incubated with endogenous mouse IgG blocking (0.13 mg/ml AffiniPure Fab Fragment (Jackson ImmunoResearch, 115-007-003) in PBS) for 1 hour at RT. Primary antibodies (Keratin 17/19 D4G2 Cell Signalling Technology 12434 and Amylase G10 Santa Cruz sc-46657) were diluted in protein block (Abcam, ab64226) and incubated overnight at 4 °C. After 3 x 5 min PBS washes, slides were then incubated with secondary antibody (1:200 in PBS) for 1 h at room temperature (AlexaFlour488 anti-mouse A28175 and AlexaFluor647 anti-rabbit, A32733). Slides were then washed 2 x PBS for 5 min, incubated with 100 μl of DAPI (1 μg/ml in PBS) for 10 min and washed again. Samples were mounted with Dako mounting media and a glass coverslip, sealed with nail varnish. Slides were imaged using a Zeiss Axioscan Z.1 at 20x magnification, images were viewed and analysed using a combination of Zeiss Zen 3.5 Blue Edition and QuPath – 0.5.0 – x64^58^.

### Cell preservation

For cryopreservation cells were seeded in T75 flask (Greiner Bio-One, 658175) for 24 h followed by trypsinisation and pelleting. Cell pellets were then resuspended in cryoprotectant consisting of the indicated percentage of DMSO (Sigma-Aldrich, D2438-50ML) and FBS. After 24 h incubation at −80 °C cells were washed 3 times with phosphate buffered saline (PBS). To remove dead cells from analyses 4′,6-diamidino-2-phenylindole (DAPI) (Sigma-Aldrich, D9542) was added to cell suspension. Cells were then moved to CellenONE (Scienion) for further processing.

For cell fixation cells were seeded in T75 flask for 24 h followed by trypsinisation and pelleting. Cell pellets were then resuspended in fixative consisting of the indicated percentage of paraformaldehyde (PFA) (Sigma, P6148) in PBS for either 20 minutes or 24 h. 20 minutes fixation was followed by tris-buffered saline (TBS) wash and subsequent PBS wash and incubation at 4 °C. 24 h incubation was performed at 4 °C. Suspension of cells were then moved to CellenONE (Scienion) for further processing.

### Pancreatic tissue processing

All animal work was approved by a University of Edinburgh internal ethics committee and was performed in accordance with institutional guidelines under license from the UK Home Office (PP7280430 and PP7510272). The mouse used for 24-hour fixation was 15 weeks old (Genotype: *Ccpg ^GT/+^*). The mouse used for 20-minute fixation was 19 weeks old (Genotype: *Pdx1-Cre;R26-LSL-CAS9-eGFP).* Pancreata were dissociated into single cells as previously described^59^. Briefly, pancreata were dissected and rinsed in 10% FBS in Hanks’ Buffered Saline Solution (HBSS, Lonza, BE10-527F) before dicing into 1-3 mm^3^ pieces on ice and centrifuging. Pancreatic fragments were enzymatically dissociated with 0.02% trypsin C-0.05% EDTA (Corning, 25-052-CI) for 10 min at 37 °C. The trypsin was inactivated with 10% FBS in DMEM and the cells washed with cold wash buffer (HBSS with 4% BSA, 0.2mg/ml soybean trypsin inhibitor (Roche, 93620-1G) and 0.2mg/ml DNase 1 (Roche, 10104159001)). A second enzymatic dissociation step was performed by suspending cells in HBSS with 4% BSA, 1.1mg/ml collagenase P (Roche, 11213857001), 0.2 mg/ml trypsin inhibitor and 0.2mg/ml DNase I and incubating for 15 min at 37 °C. The tissue was then mechanically dissociated by serological pipetting. Light microscopy was used to assess cell dissociation and viability (with trypan blue to mark dead cells). Incubation was continued until 90% of cells were single cells or when a reduction in viability was detected. Cells were washed twice with HBSS containing 5% FBS and 0.2mg/ml DNase I, passed through a 70µm nylon mesh and resuspended in 2ml cold wash buffer. All centrifuge steps in the protocol were performed at 350 x g at 4 °C for 5 minutes. Live cells were sorted for analysis using a BD LSR Fortessa X-20 analyser, gating out dead cells stained with DAPI.

### Cell sample processing for mass spectrometry

Single cell seeding and processing was performed as previously described^20,60^. Briefly, 300nL of master mix (0.2% DDM (Sigma-Aldrich, D4641-5G), 100 mM TEAB (Supleco, 18597), 10 ng/µL trypsin (Thermo Scientific, 90058)) was dispensed into proteoCHIP EVO 96 plate (Scienion, C-PEVO-96-BUN) or 384 well plate (Eppendorf, 0030129547) inside the CellenONE. Cells matching the isolation criteria were dispensed to each well and incubated at 50°C with 85% relative humidity for 1.5 hours within the instrument. Post incubation, the temperature was reduced to 20°C and 3.2 µL of 0.1 % formic acid (FA) was added to the samples. Subsequently, samples were loaded onto purification and loading trap columns (Evosep, Evotips). Evotips were prepared according to manufacturer’s instructions. The samples were loaded onto the Evotips by centrifugation at 800 g for 60 seconds (from proteoCHIP EVO 96 plate) or manually (from 384 well plate). After loading, the Evotips are washed with 0.1% FA followed by a final wash of 100 µL 0.1% FA and spinning for 10 seconds at 800 g. The samples are then transferred into Evosep One LC system for LC-MS/MS analyses.

### LC-MS/MS

LC-MS/MS analyses were performed on timsTOF SCP Mass Spectrometer (Bruker) coupled to Evosep One LC system (Evosep). We utilised Aurora Elite CSI analytical columns (IonOpticks, AUR3-15075C18-CSI) and CaptiveSpray ionization source. Samples were analysed using the Whisper 40 SPD method (gradient flow of 100 nl/min with a 31 minutes method duration) or Whisper zoom 40 SPD (gradient flow of 200 nl/min with a 31 minutes method duration). Eluted peptides were analysed with a parallel accumulation-serial fragmentation data-independent acquisition (diaPASEF) method. Isolation windows optimised for low input samples using open-source Python package for dia-PASEF methods with Automated Isolation Design (py_diAID)^61^. Two methods were used for data acquisitions: a window placement scheme consisting of 8 TIMS ramps with 3 mass ranges per ramp spanning from 400 – 1000 m/z and from 0.64 – 1.40 1/K0; a window placement scheme consisting of 4 TIMS ramps with 5 mass ranges per ramp spanning from 327 – 1200 m/z and from 0.7 – 1.30 1/K0. Spectronaut 18 was used for data analysis searching against Uniprot *Homo sapiens*, *Mus musculus* or *Escherichia coli* database. Readout from each search can be found in Supplementary Table 1.

For entrapment analysis, *Escherichia coli* the fasta file distributed by Spectronaut (uniport_sprot_2024-01-01_ECOLI, 4530 entries) was used. Trypsin/P was specified as protease and one missed cleavage was allowed. Precursor FDR was set to the indicated percentage.

### Data analyses

Analysis of proteomic data was performed using R version 4.3.3. R/Bioconductor package MSnbase was used for missing values imputation using MinProb method^62,63^. Outliers were removed from datasets. UMAP, PCA and PLS-DA plots were created using R packages: dplyr (version 1.1.4), UMAP (version 0.2.10.0), ggplot2 (version 3.5.1), mixOmics (version 6.26.0). Further statistical calculations, such as Student’s *t*-tests, ANOVA and z-score were calculated using Perseus software^64^. Heatmap and hierarchical clustering was conducted using Morpheus online tool (https://software.broadinstitute.org/morpheus). Volcano plots were created using VolcaNoseR online tool^65^. Gene ontology analyses, corresponding heatmaps and Circos plots were obtained using Metascape and DAVID online tools^66,67^. All illustrations were created with BioRender.com.

## Acknowledgement

All illustrations were created with BioRender.com. We thank Alejandro Brenes (University of Edinburgh) for helpful discussions. We are grateful to members of the von Kriegsheim laboratory for discussions and critical reading of the manuscript. S.W is supported by CRUK Senior Fellowship (C20685/A29576); A.V.K. is supported by MRC fellowship (MR/X01293X/1) and BBSRC fellowship (BB/X019160/1).

## Author contributions

Conceptualisation, A.N.M. and A.V.K.; Methodology, A.N.M., J.H., J.M., S.W and A.V.K.; Investigation, A.N.M., J.H., J.M., S.I., S.W and A.V.K.; Writing – Original Draft, A.N.M. and A.V.K.; Writing – Review and Editing, all authors; Funding Acquisition, A.V.K.

