## Supplementary Figures for "Overcoming preservation challenges to enable single-cell proteomics of fixed cell and tissue samples with retained proteome integrity"

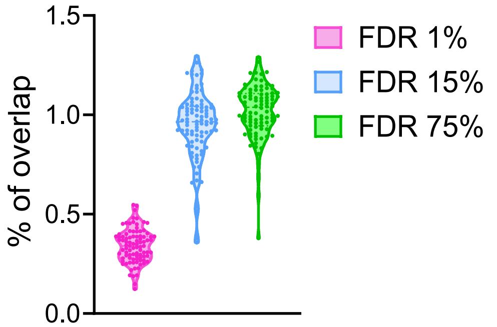


**Supplementary Figure 1 (related to Figure 1). Entrapment sequence method using E.coli library.**

Violin plot showing percentage of identified E.coli proteins in the extended human database search at stated FDR settings.

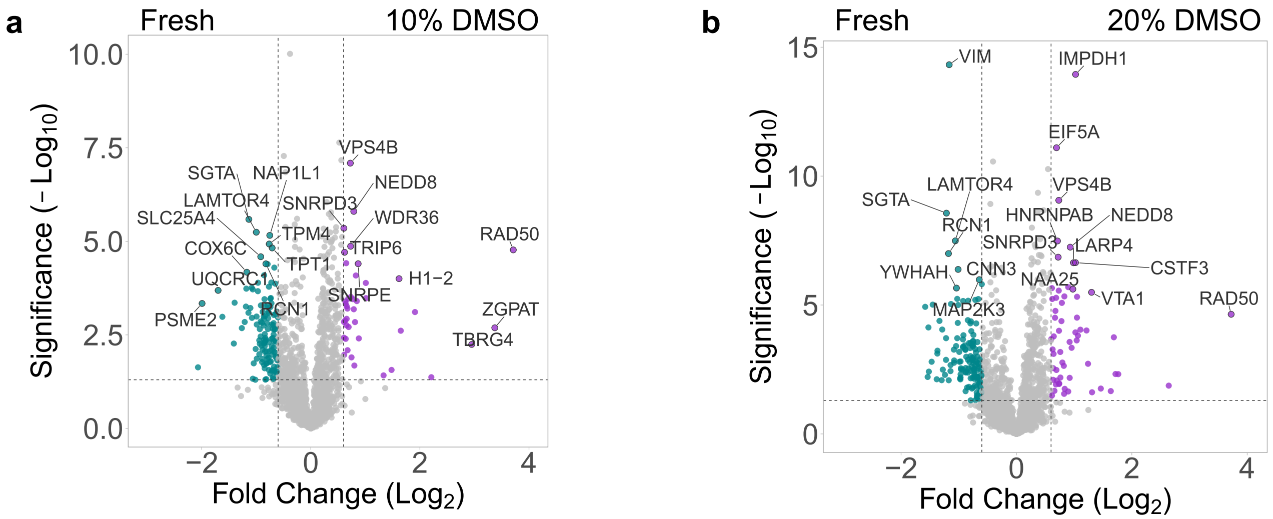


**Supplementary Figure 2 (related to Figure 2). Direct comparison of tested preservation method to fresh cell proteome.**

(a) Volcano plot of two-sample t-test results (10% DMSO over Fresh). Log2 fold change and −log10 p-value are shown, significantly regulated proteins are shown in cyan and purple.

(b) Volcano plot of two-sample t-test results (20% DMSO over Fresh). Log2 fold change and −log10 p-value are shown, significantly regulated proteins are shown in cyan and purple.

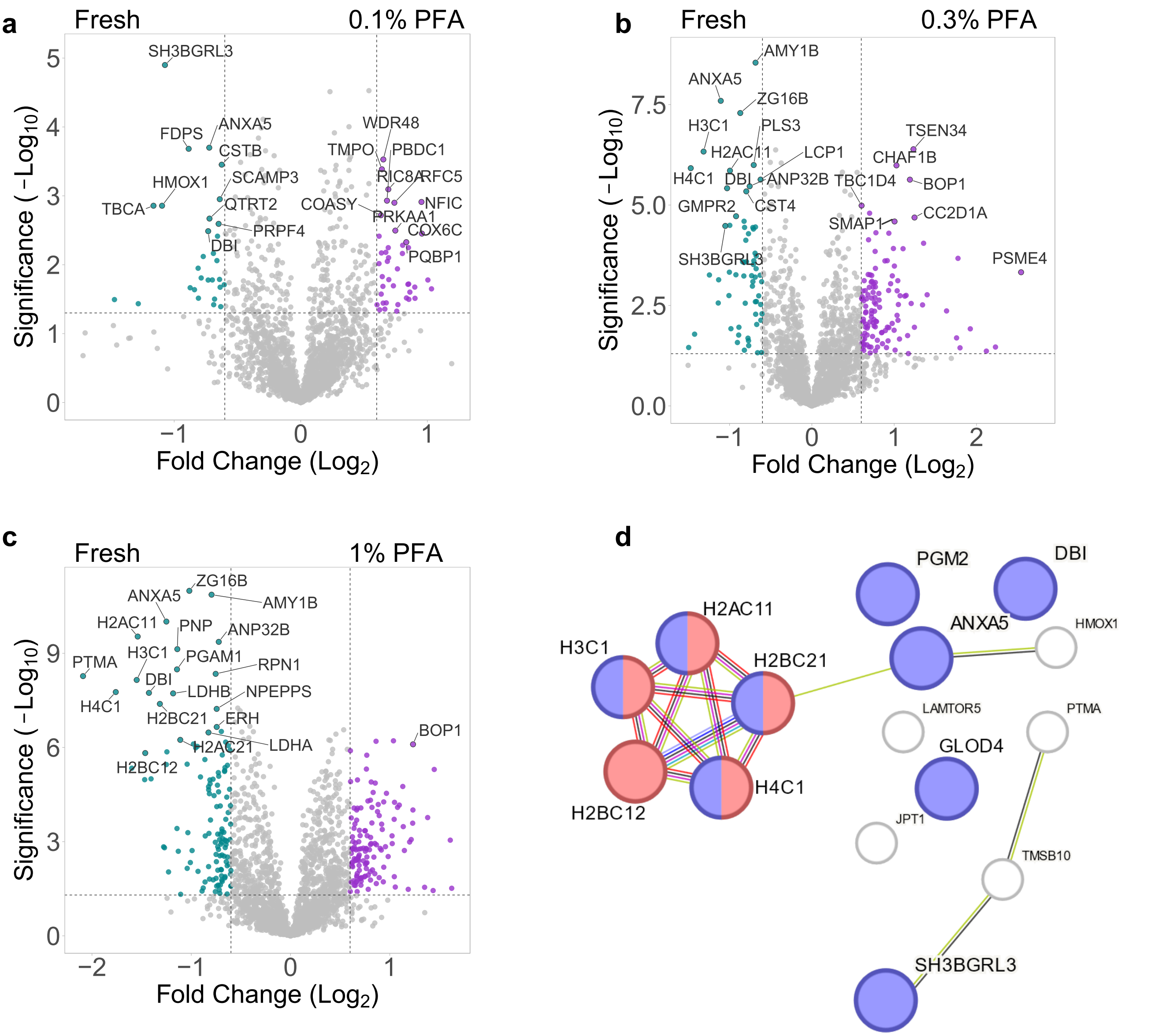


**Supplementary Figure 3 (related to Figure 3). Direct comparison of tested preservation method to fresh cell proteome.**

(a) Volcano plot of two-sample *t*-test results (0.1% PFA over Fresh). $Log_2$ fold change and $-log_{10}$ p-value are shown, significantly regulated proteins are shown in cyan and purple.

(b) Volcano plot of two-sample *t*-test results (0.3% PFA over Fresh). $Log_2$ fold change and $-log_{10}$ p-value are shown, significantly regulated proteins are shown in cyan and purple.

(c) Volcano plot of two-sample *t*-test results (1% PFA over Fresh). $Log_2$ fold change and $-log_{10}$ p-value are shown, significantly regulated proteins are shown in cyan and purple.

(d) String analysis of top 15 hits significantly decreased in (c) based on fold change, histone proteins are highlighted in red, exosome proteins are highlighted in blue.

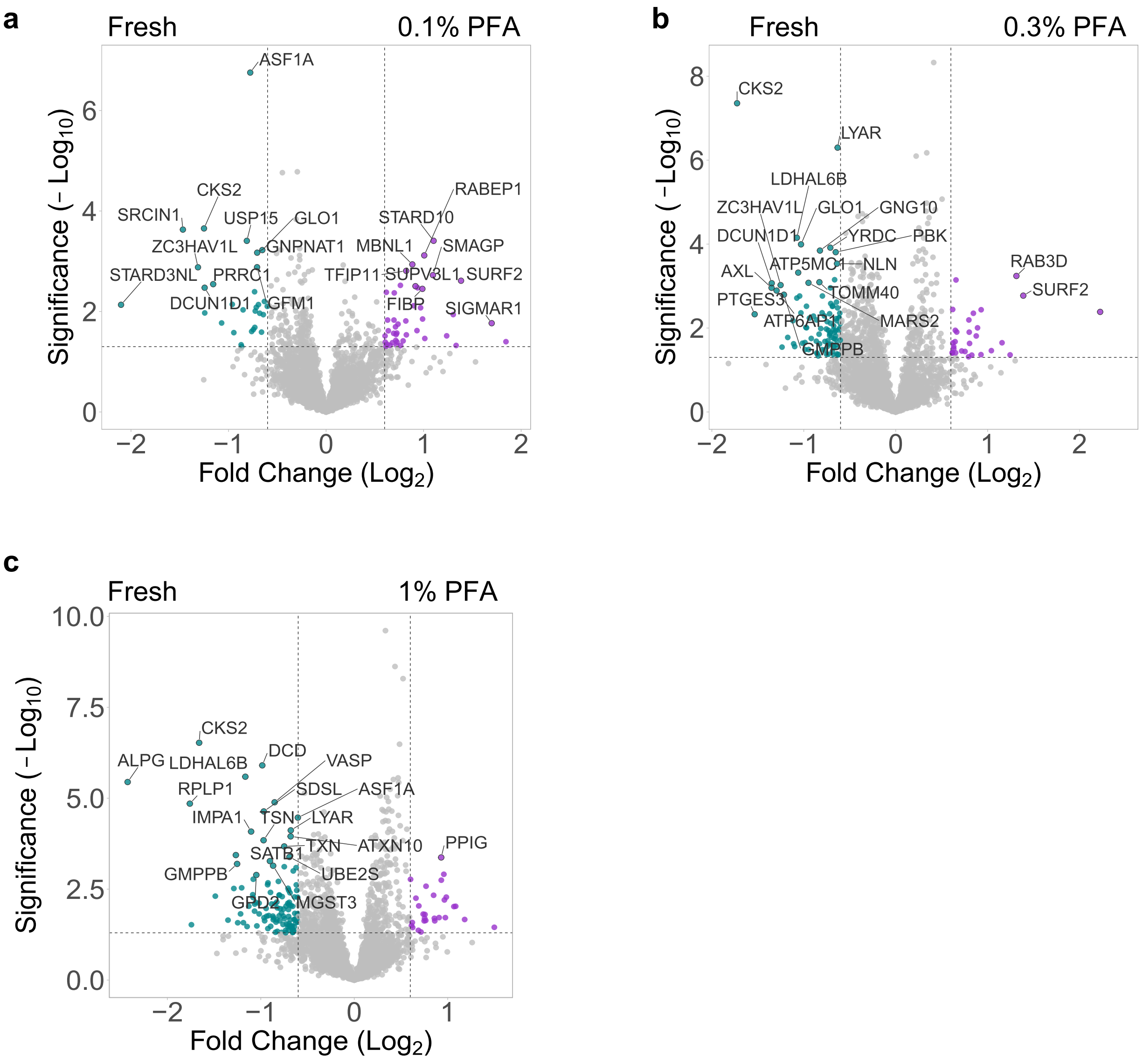


**Supplementary Figure 4 (related to Figure 4). Direct comparison of tested preservation method to fresh cell proteome.**

(a) Volcano plot of two-sample *t*-test results (0.1% PFA over Fresh). $Log_2$ fold change and $-log_{10}$ p-value are shown, significantly regulated proteins are shown in cyan and purple.

(b) Volcano plot of two-sample *t*-test results (0.3% PFA over Fresh). $Log_2$ fold change and $-log_{10}$ p-value are shown, significantly regulated proteins are shown in cyan and purple.

(c) Volcano plot of two-sample *t*-test results (1% PFA over Fresh). $Log_2$ fold change and $-log_{10}$ p-value are shown, significantly regulated proteins are shown in cyan and purple.

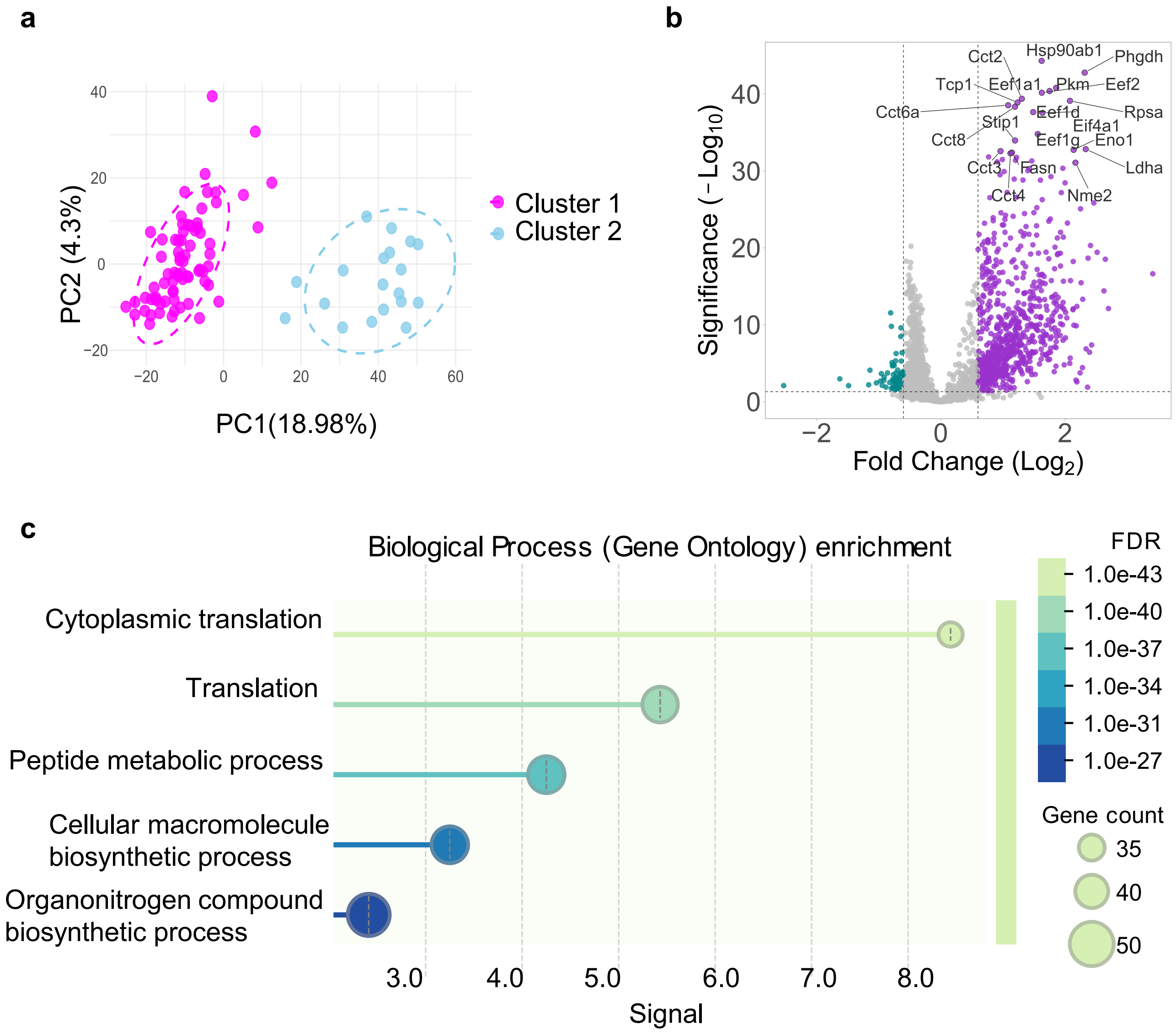


**Supplementary Figure 5 (related to Figure 5). Proteomics analyses of PDAC single-cell samples.**

(a) PCA plot of PDAC single-cell sample group: Cluster 1 (magenta) and Cluster 2 (blue).

(b) Volcano plot of two-sample *t*-test results (Cluster 2 over Cluster 1) of PDAC single-cell proteomics analyses. $Log_2$ fold change and $-log_{10}$ p-value are shown, significantly regulated proteins are shown in cyan and purple.

(c) Functional enrichment visualisation of the top 5 GO biological processes enriched in Cluster 2 of PDAC single-cell dataset. Size of the dot corresponds to gene number of the indicated biological process. The gene count dots are coloured by their false discovery rate (FDR).

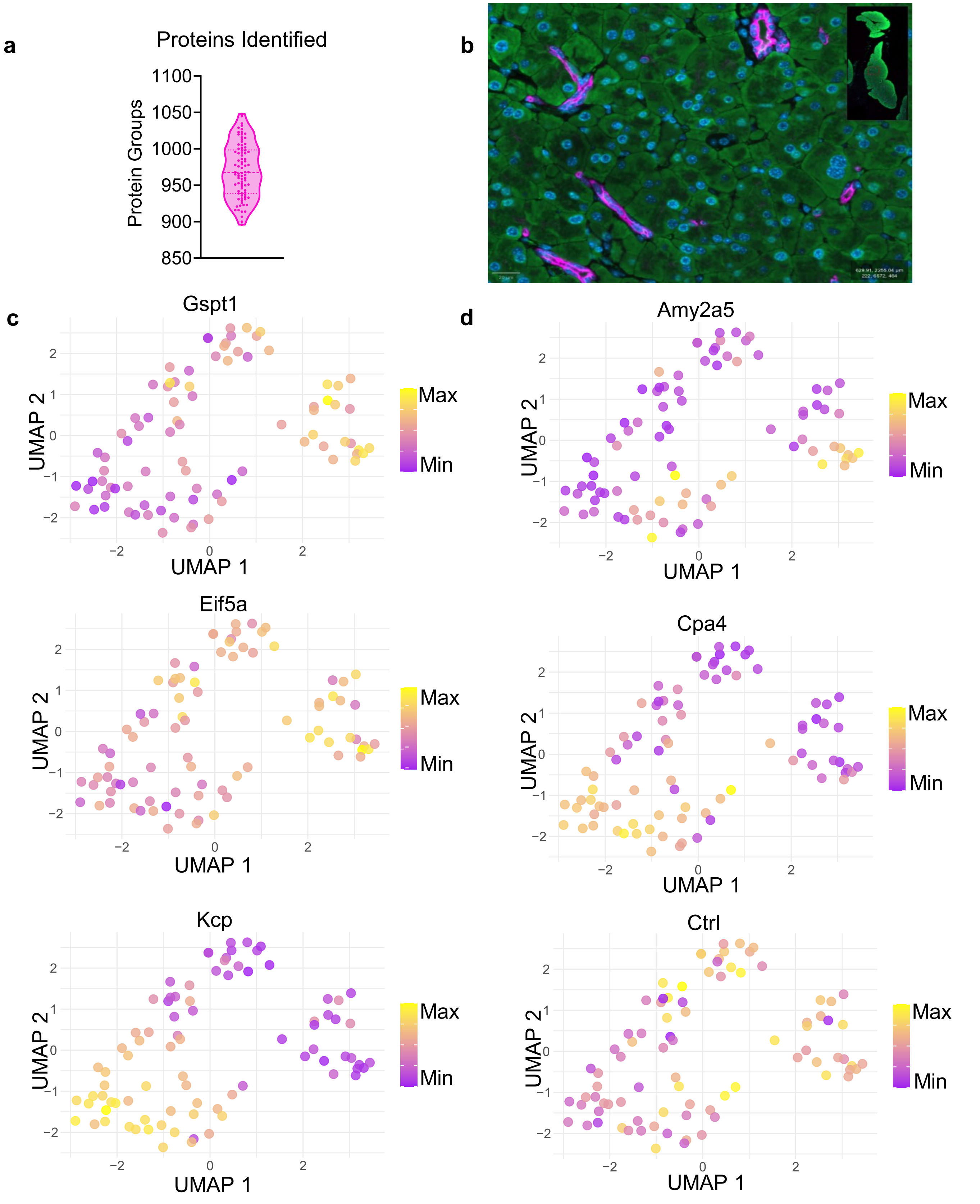


**Supplementary Figure 6 (related to Figure 7). Proteomics analyses of single-cell derived from mouse pancreas tissue.**

(a) Violin plot showing protein group identifications from single-cells pancreatic tissue sample. Each dot represents readout from a single-cell.

(b) A representative immunofluorescence image of mouse pancreas with exclusive positive amylase staining of acinar cells (green) and cytokeratin 17/19 staining of ductal cells (magenta). Nuclei are stained with DAPI (blue). Scale bar 20um.

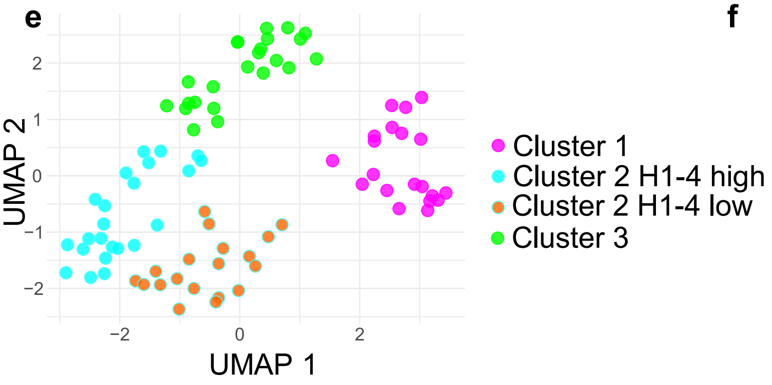


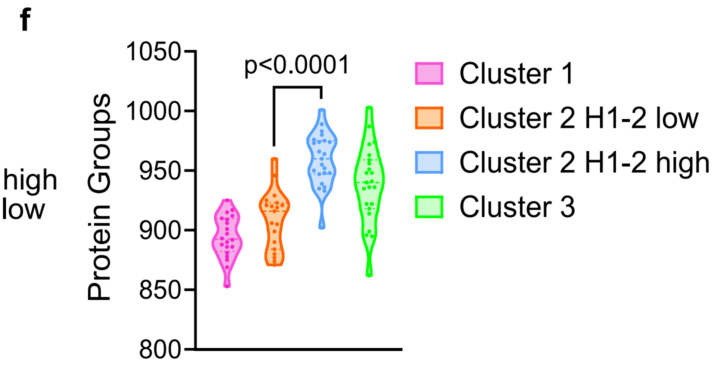


**Supplementary Figure 6 (related to Figure 7). Proteomics analyses of single-cell derived from mouse pancreas tissue cont.**

(c) UMAP plot of acinar tissue single-cell dataset, overlaid with absolute values of selected proteins detected in the dataset. The lowest value in the dataset is represented in purple (Min), the highest Gspt1 value in the dataset is represented in yellow (Max).

(d) UMAP plots of acinar tissue single-cell dataset, overlaid with absolute values of selected pancreatic enzymes. The lowest value in the dataset is represented in purple (Min), the highest value in the dataset is represented in yellow (Max).

(e) UMAP of samples groups with separation of Cluster 2 based on H1-4 levels. Cluster 1 (magenta), Cluster 2 H1-4 low (blue), Cluster 2 H1-4 high (orange), Cluster 3 (green)

(f) Violin plot showing protein group identifications from single-cells as seen in (a). Each dot represents readout from a single-cell. P-values were calculated using an unpaired, two-tailed t-test.
